# Optimization of an unscented Kalman filter for an embedded platform

**DOI:** 10.1101/2022.01.11.475885

**Authors:** Philip P. Graybill, Bruce J. Gluckman, Mehdi Kiani

## Abstract

The unscented Kalman filter (UKF) is finding increased application in biological fields. While realizing a complex UKF system in a low-power embedded platform offers many potential benefits including wearability, it also poses significant design challenges. Here we present a method for optimizing a UKF system for realization in an embedded platform. The method seeks to minimize both computation time and error in UKF state reconstruction and forecasting. As a case study, we applied the method to a model for the rat sleep-wake regulatory system in which 432 variants of the UKF over six different variables are considered. The optimization method is divided into three stages that assess computation time, state forecast error, and state reconstruction error. We apply a cost function to variants that pass all three stages to identify a variant that computes 27 times faster than the reference variant and maintains required levels of state estimation and forecasting accuracy. We draw the following insights: 1) process noise provides leeway for simplifying the model and its integration in ways that speed computation time while maintaining state forecasting accuracy, 2) the assimilation of observed data during the UKF correction step provides leeway for simplifying the UKF structure in ways that speed computation time while maintaining state reconstruction accuracy, and 3) the optimization process can be accelerated by decoupling variables that directly impact the underlying model from variables that impact the UKF structure.

**Author summary:** Powerful statistical tools can be used to estimate unmeasured information in biological systems and predict the system’s future state. Translating these tools from a desktop computer to small, wearable or implantable platforms can unlock many benefits, but presents many design challenges, because tradeoffs between computational simplicity and system accuracy must be balanced. We present an approach to translating one such widely used statistical tool, the unscented Kalman filter, from a desktop computer to a miniature device. We demonstrate the approach with a case study centering on the neural circuits governing the mammalian sleep-wake cycle. By exploiting uncertainty in the biology being estimated and the corrective influence of biological measurements, we arrive at a design that is easily computable on the miniature device and maintains a high level of accuracy. We anticipate that our approach will aid others in designing such miniaturized systems for a broad range of applications.

## Introduction

Nonlinear Kalman filtering presents exciting possibilities for research in dynamical systems as applied to biology. Among nonlinear Kalman filters, the unscented Kalman filter (UKF) has gained widespread use due to its excellent performance and its deterministic, gradient-free implementation. Through the UKF, physiological states that are difficult or costly to measure can be estimated by the assimilation of more easily acquired or less costly observables [1]. Model states and parameters can be tracked to gain insight into physiological phenomena. And future state forecasts can be made to guide intervention.

Two early studies that demonstrated the utility of the UKF in biological systems employed computer simulations. Voss *et al*. showed that a single state of model-generated data with added noise could be used to reconstruct unmeasured states in a FitzHugh-Nagumo neuron model [2]. In [3], UKF state and parameter estimates were used in closed-loop control of potential waves in a computational model of cerebral cortex.

More recent studies have used the UKF to assimilate *in vivo* or *in vitro* data in physiological models. In [4], patient data were used to estimate states and parameters in models of the human respiratory system. In [5], *in vitro* neuron membrane potential recordings were assimilated in a model of a small network of Hodgkin-Huxley-type neurons. Other studies have applied the UKF to estimate parameters in the Cad system in E. coli [6], track anesthetic brain states using patient EEG [7], and forecast blood glucose levels for diabetic patients using individually-tailored models [8]. Bahari *et al*. employed a UKF to predict sleep-state transitions using *in vivo* neural recordings from chronically implanted animals [9].

Deploying a UKF or related nonlinear Kalman filter (KF) in an embedded system provides numerous advantages over traditional computational platforms in the context of biological systems. The use of embedded hardware dedicated to the specific application allows for tight control over task scheduling, minimizes feedback latency in closed-loop applications, and avoids interruptions in communication that can occur with more complex operating systems. Embedded systems can often be converted into a wearable form. Wearability aids in the constant monitoring of biological signals, increases freedom of movement for the subject or animal, and typically simplifies use and maintenance, which may increase the likelihood of long-term user adoption. For biomedical devices used in translational animal experiments, wearability may ease the path from animal trials to human trials. The embedded deployment of UKFs is also relevant in certain non-biological contexts, such as navigation systems in small unmanned aerial vehicles (UAVs).

However, realization of a linear or nonlinear KF in an embedded system can prove challenging. Embedded processors have limited computational speed compared to their non-embedded counterparts and typically have only one core. In wireless embedded systems, power constraints related to battery size or wireless power transfer may further limit computation speed. In a closed-loop application, the required rate of feedback delivered to the system sets a real-time processing deadline. In the interval between subsequent instances of feedback, the CPU must perform any communication tasks, integrate the model equations forward in time, and evaluate the UKF equations, which include matrix operations. The feedback signal may depend upon the predicted system state at some forecast horizon time beyond the next scheduled feedback. This forecasting extends the time required to integrate the model equations. In short, the limitations of the embedded platform and computational demands of the UKF can combine to render the meeting of the real-time requirement impossible for even a relatively small state-space dimension.

Several efforts have been made to deploy linear and unscented KFs in embedded systems. In [10], a UKF for nanosatellite attitude determination was implemented on a field-programmable gate array (FPGA). The same authors also designed an FPGA-microcontroller (MCU) system that included an FPGA block that performed the generic UKF computations and a flexible software implementation of the model-specific computations [11, 12]. In [13], the authors implemented a UKF for a navigation system on an MCU and minimized execution time by utilizing processor-specific matrix libraries, replacing calls to functions with in-line code, changing constant divisions to multiplications by the constants’ inverses, and testing different MCUs. A systematic approach for reducing computation time in embedded systems utilizing the KF, UKF, and the extended KF was outlined in [14]. In [15], a KF was implemented on an FPGA for decoding rat paw movement from neural spike data. While each of these works presents a sufficient system for real-time usage and a subset of them reduce computation time by exploring the design space, none of them provide a comprehensive consideration of potential optimization parameters for reducing a UKF system for embedded deployment or a method for actual optimization through minimization of cost functions.

Here, we present a method for optimizing the design of a UKF system for deployment on a given MCU or soft processor in FPGA fabric. The method is generalizable to any model. The optimization searches over a large set of variants in both the formulation of the underlying model and the structure of the UKF itself. Furthermore, we apply a multi-objective cost function that accounts for the competing objectives of minimizing computation time and power consumption and minimizing error in UKF estimation and forecasting. To demonstrate the optimization method, we present a case study using data generated from a model of the sleep-wake regulatory network [16]. We leave iteration of the method across computing platforms to the reader.

In addition to offering a generalized approach to fitting a UKF on an embedded platform, we highlight three main insights from our work: 1) The magnitude of the model process noise can be leveraged to justify using simpler, less accurate integration schemes and model approximations. 2) The constraining influence of the UKF data assimilation step, or measurement correction, may permit the use of a computationally simpler UKF structure without significantly sacrificing accuracy in state estimation. 3) And the UKF optimization can be accelerated by thoughtfully ordering the optimization stages and separating the swept variables into two classes—model variables and UKF structure variables. By eliminating unsuitable combinations of model variables through forecast integration tests, the overall optimization space can be reduced before more time-intensive UKF state reconstruction tests are performed.

In the “Materials and methods” section, we review the UKF formalism to establish notation and draw attention to the aspects of the UKF that undergo variation in the optimization. We also introduce the Diniz Behn and Booth (DBB) model of the rat sleep-wake regulatory network, which we use for our case study. The target MCU platform and a proxy computation platform used to compute the bulk of the optimization data are also described. We open the “Results” section with a description of the variable space over which the optimization is run. We then detail the optimization method stage by stage and weave in discussion of the case study results in order to illustrate the method. In the “Conclusion” we summarize the method, highlight our insights, and propose future work.

## Materials and methods

Our case study scenario for the UKF optimization method is a real-time *in vivo* sleep-wake regulation experiment in rats, in which feedback to the animal is determined by 10-second forecasts of future state. Our computational target is a system-on-chip/FPGA (SoC-FPGA) combining an MCU and FPGA fabric. We aim to deploy the UKF on the MCU only, and we reserve the FPGA fabric for signal processing of multiple input channels. Here, we review relevant computational details of the UKF, the DBB model of sleep-wake regulation that provides the kernel for the UKF, and details of the target and proxy computing platforms employed in the optimization.

### UKF formalism

Sigma point filters are a family of adaptations of the linear Kalman filter for nonlinear systems. In sigma point filters, the distribution of state is represented through a set of deterministically chosen “sigma points.” Each sigma point is propagated forward in time through the nonlinear dynamics to the next scheduled measurement of data. The forward-iterated sigma points are then used to form a linearization of the dynamics through which the measured data are assimilated to the estimated distribution. We use the term “unscented Kalman filter” to refer to any of a number of sigma point filters that employ some version of the unscented transform for the selection of sigma points [17–21]. We depict the UKF graphically in Fig 1A. The formalism presented here follows that in [22] and [23].

**Fig 1.**
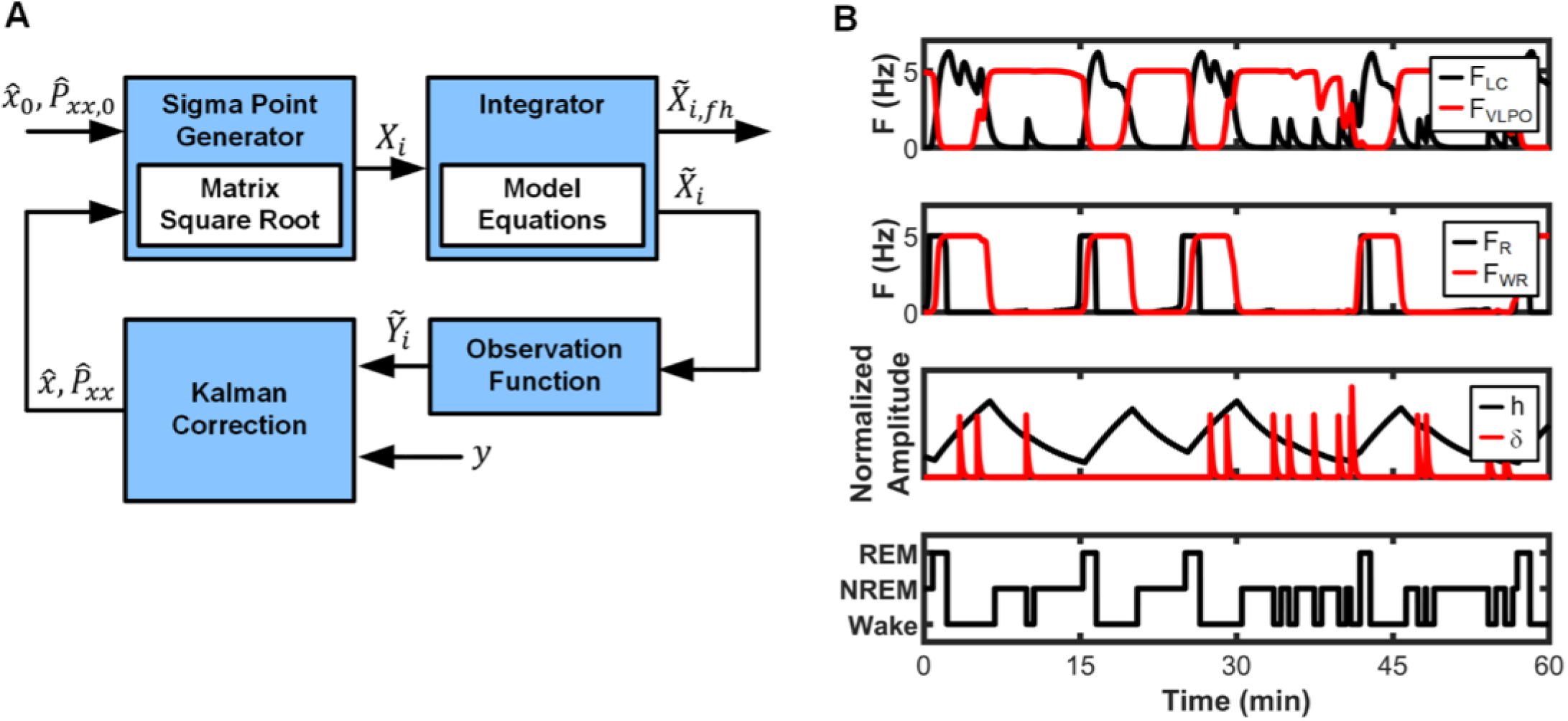
UKF block diagram and Diniz Behn and Booth (DBB) model output. (A) The unscented Kalman filter consists of a repeating cycle of two phases: 1) prediction of state distribution, in which representative sigma points *X*_*i*_ are generated and propagated through the nonlinear model equations, and 2) correction, in which observed data *y* is assimilated to the estimated distribution. The sigma points can be integrated beyond the next scheduled observation 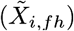 to produce a forecast of state distribution. (B) Example DBB model states, including mean population firing rates for the wake-active LC, NREM-active VLPO, and REM-active and wake-REM-active subpopulations of the LDT/PPT. Normalized homeostatic sleep pressure *h*, normalized noisy input *δ*, and derived hypnogram are shown in the bottom two panels.

The discrete-time equation

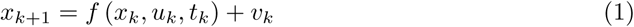

models the dynamics of an *n*-dimensional system, where *u*_*k*_ is system input, *t*_*k*_ is the discretized time at step *k*, and *v*_*k*_ is the process noise. The measurement of an *m*-dimensional variable *y* is described by the linear or nonlinear observation function

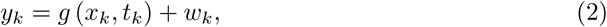

where *w*_*k*_ is the observation noise. The noise terms *v*_*k*_ and *w*_*k*_ are uncorrelated, zero-mean Gaussian vectors with covariance matrices *Q*_*k*_ (*n* × *n*) and *R*_*k*_ (*m* × *m*), respectively. The process noise accounts for aspects of the true system that the model *f* (·) fails to capture, and the observation noise accounts for uncertainty in the measurement.

Like other Kalman filters, The UKF consists of alternating steps of *prediction* and *correction* to produce an estimate of state distribution with mean 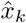 and covariance 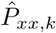. At the start of every prediction-correction cycle, a set of sigma points *X*_*i*_, *i* = 0, …, *N*, is chosen to represent the estimated state distribution by satisfying the constraints

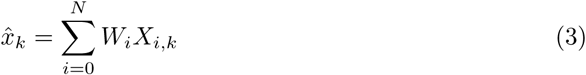

and

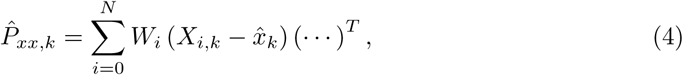

where *W*_*i*_ are scalar weights, and (· · ·) is shorthand for the preceding term in the equation. In UKFs, the computation of the sigma points *X*_*i,k*_ requires a square root of the matrix 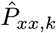. In the *prediction* step, the sigma points are propagated through the nonlinear dynamics in Eq (1). The predicted mean 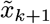 and predicted covariance 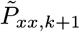 are found from the forward-iterated sigma points:

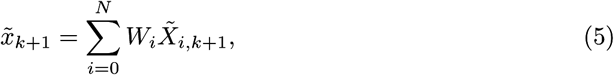

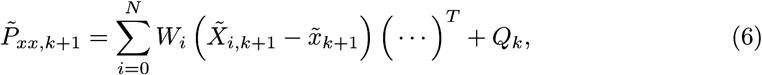

where 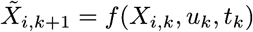, and the tilde denotes predicted data that have not yet undergone the correction step.

Next, the forward-iterated sigma points 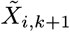 are transformed into the observation space using Eq (2): 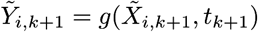.^1^ The 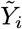’s are used to find the predicted mean and covariance transformed into the observation space:

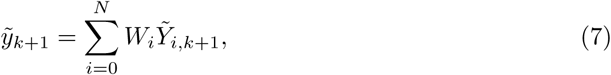

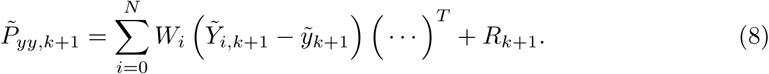

In the *correction* step, the Kalman gain is found:

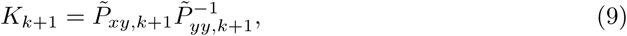

where the cross-covariance is defined as

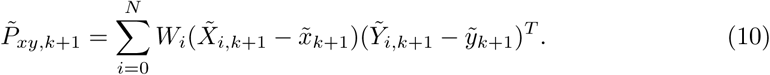

The observed data *y*_*k*+1_ are then used to find the corrected mean,

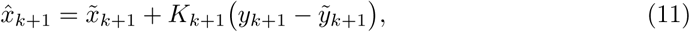

and the corrected covariance is found from the Kalman gain and 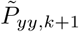,

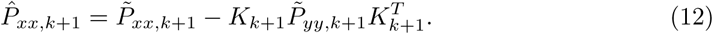

The hat indicates terms that have undergone the correction step. The corrected mean and covariance are then used to produce the sigma points for the next cycle, *X*_*i,k*+1_.

Numerical integration is used to advance the sigma points from *t*_*k*_ to *t*_*k*+1_ using the dynamics in Eq (1). However, the integrator can also be used to forecast the state well beyond the next anticipated observation time by integrating the sigma points to a given forecast horizon. Therefore, although the integrator is not an explicit part of the UKF equations, its implementation impacts estimation and forecast accuracy as well as computation time. In order to avoid confusion, we refer to the computation of 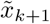 and 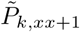 as *prediction* and to the integration of the state equations beyond the next anticipated observation as *forecasting*.

### The Diniz Behn and Booth model

As a case study, we ran our optimization procedure on a UKF system for forecasting neural activity in a behaving rat. We used Diniz Behn and Booth’s model of the sleep-wake regulatory system [16] as the model in Eq (1). In the Diniz Behn and Booth (DBB) model, whose twelve states are listed in Table 1, the relative firing rates of five neural populations *F*_*z*_ indicate the animal’s state as wakefulness (W), rapid-eye-movement (REM) sleep, or non-REM (NREM) sleep. The five populations are the locus coeruleus (LC), dorsal raphe (DR), ventrolateral preoptic nucleus (VLPO), a REM-promoting subpopulation from the laterodorsal tegmentum (LDT) and pedunculopontine tegmentum (PPT), and a wake/REM-promoting subpopulation from LDT and PPT. Interactions between the neural populations are mediated by the normalized concentrations of the neurotransmitters *C*_*i*_ emitted by each of the five populations. The two other model states are the homeostatic sleep pressure variable *h* and the random, wake-inducing input to the LC and DR, called *δ*. Example outputs for select DBB model states are shown in Fig 1B.

**Table 1.** States in the Diniz Behn and Booth model.

| No. | State | Abbrev. |
| --- | --- | --- |
| **Population average firing rates** |  |  |
| 1 | Locus coeruleus (LC) (wake active) | $F_{LC}$ |
| 2 | Dorsal raphe (DR) (wake active) | $F_{DR}$ |
| 3 | Ventrolateral preoptic nucleus (VLPO) (NREM active) | $F_{VLPO}$ |
| 4 | REM promoting subpopulations in the laterodorsal and pedunculopontine tegmenta (R) | $F_R$ |
| 5 | Wake/REM promoting subpopulations in the laterodorsal and pendunculopontine tegmenta (WR) | $F_{WR}$ |
| **Normalized neurotransmitter concentrations** |  |  |
| 6 | Noradrenaline | $C_N$ |
| 7 | Serotonin | $C_S$ |
| 8 | γ-aminobutyric acid (GABA) | $C_G$ |
| 9 | Acetylcholine from the R subpopulation | $C_{AR}$ |
| 10 | Acetylcholine from the WR subpopulation | $C_{AWR}$ |
| **$h$ and $\delta$** |  |  |
| 11 | Homeostatic sleep drive | $h$ |
| 12 | Noisy input to the LC and DR | $\delta$ |

The five average firing rates are governed by the equations

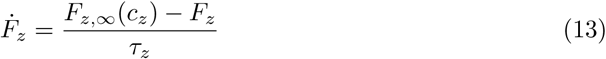

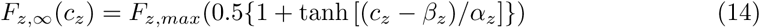

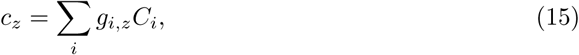

where the subscript *z* ∈ {*LC, DR, V LPO, R, WR*} denotes one of the five neural populations; *F*_*z*,∞_ is the steady-state firing rate for population *z*; *c*_*z*_ is the summed excitatory and inhibitory neurotransmitter input to population *z*; and *τ*_*z*_ governs the rate of decay from *F*_*z*_ to *F*_*z*,∞_. The tanh function in Eq (14) serves to as a smooth step function between near-zero and maximum firing rates, and the parameters *β*_*z*_ and *α*_*z*_ govern the midpoint and steepness of the step, respectively. The constants *g*_*i,z*_ define the degree of excitatory or inhibitory effect of neurotransmitter *i* on population *z*. The *g*_*i,z*_ parameters should not be confused with the observation function *g*(*x*_*k*_) in Eq (2).

Each model population emits a specific neurotransmitter. LC emits noradrenaline, DR emits serotonin, VLPO emits GABA, and the subpopulations from LDT/PPT both emit acetylcholine. The equations for the *C*_*i*_’s have roughly the same form as the *F*_*z*_’s, but take as input only the firing rate of a single population:

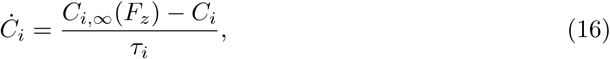

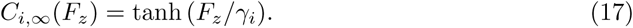

The result of Eq (17) is multiplied by a noise factor drawn from a Poisson process in order to model the variability of neurotransmitter output in relation to the population firing rate.

The homeostatic sleep drive *h* increases while the model animal is awake and decreases while the animal is asleep. Wake/sleep state is determined by summing the firing rates of the wake-active populations *F*_*LC*_ and *F*_*DR*_ and comparing the sum with the wake threshold parameter *θ*_*W*_ :

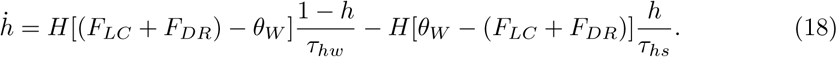

The variable *h* enters the formula for *F*_*V LP O*,∞_ through the parameter *β*_*V LP O*_(*h*) = −*kh*.

Finally, noisy input to LC and DR is modeled as a train of randomly occuring impulses *δ*_*in*_ with varying amplitude and a decay rate governed by the time constant *τ*_*δ*_:

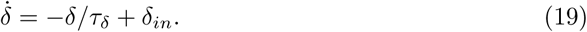

We point out two qualities of the DBB model that impact computation in real-time embedded systems and are also common to other biological models. One, the tanh function appears multiple times in the model equations. Sigmoid functions such as tanh are commonly used in neural mass models as transfer functions from population input to population output, as well as in other biological models in which a state transitions smoothly from a minimum “off” condition to a maximum “on” condition. Such sigmoid functions can account for a large portion of the operations in an embedded model integrator, and substituting an approximation for the sigmoid can yield significant reduction in computation time. Two, random biological noise relaxes the required level of fidelity of the numerical integrator to the ideal model equations. In application, the noisy input to the system *δ*(*t*) is practically unknown. Therefore, when the model equations are integrated, the input is set to zero, and the model noise *Q* is increased accordingly to account for the uncertainty of input. Although a larger *Q* further spreads the estimated distribution of state, it permits the use of a computationally simpler integrator. As long as the deviation of a simplified integrator from a “best available” integrator is small with respect to the noise *Q*, the simplified integrator can be safely substituted for one with higher fidelity to the ideal model without significantly impacting the forecast accuracy.

### Computation platforms

The amount of time required to process all of the optimization data for the tested variants on the target MCU is prohibitive, due to the large quantity of test data and the limited computing speed of the MCU. For this reason, we used a desktop PC as a proxy platform for computing the larger volumes of optimization data, and we reserved the target MCU platform for only measuring the time required to compute one cycle of each of the tested UKF variants. The target platform for the case study was a 32-bit Arm Cortex-M3 processor contained in a SmartFusion2 SoC-FPGA (M2S010, Microsemi, Aliso Viejo, CA, USA) running at 100 MHz. The Cortex-M3 does not include a floating point unit. MATLAB (The MathWorks, Natick, MA, USA) running on a desktop computer with a 64-bit Intel Core i7-6700 CPU (3.4 GHz) served as the proxy computation platform. All UKF variants under test were programmed in MATLAB, and these proxy variants were used to compute the data for the stages of the optimization in which forecast divergence and state reconstruction accuracy were tested.

## Results

We present here a method for the design of an embedded UKF system as a true multi-objective optimization process—in the sense of minimization of a cost function—that balances sufficiently fast computation time and low power on the one hand and sufficient fidelity of state forecasting and estimation on the other hand. Here, *fidelity* refers to how closely a computed test output matches a target benchmark output, which may or may not closely approximate the true state of the system. The goal of the optimization is to find an alternative embedded computation scheme that has high fidelity relative to the more expensive benchmark scheme. The error between the benchmark and the true system state is taken as a reference point for assessing the fidelity of the alternative.

The optimization is divided into three test stages, which are mapped in Fig 2: 1) a *computation time* test over all variants, 2) a *forecast* test over remaining *model* variants only, and 3) a *state reconstruction* test over all remaining variants. Each of the stages produces a scalar result for each tested variant, which is compared to a threshold value. Combinations of variables producing a result below the threshold pass to the next stage, while those exceeding the threshold are excluded from further testing. Variants that pass the third stage are assessed with a cost function that combines the scalar results of all three stages. The *benchmark variant* is a version of the UKF that has been developed on a desktop PC to yield excellent state reconstruction fidelity with respect to the true system, but which has been developed without being constrained by the computational limitations of the target MCU. Performance of all other computational variants are compared to this benchmark.

**Fig 2.**
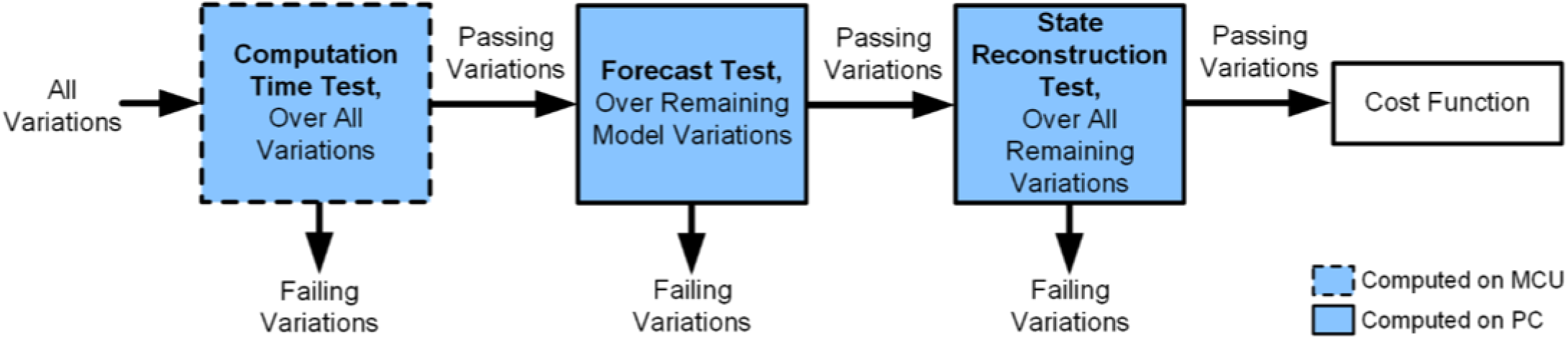
Optimization stages. The optimization comprises three main test stages: computation time test, forecast test, and state reconstruction test. Variants passing all three stages are assessed with a cost function. Only the computation time test is computed on the target MCU.

### Optimization variable space

We divide UKF design variables into three broad classes—fixed variables, model variables, and UKF structure variables—and identify the specific values we allowed them to take in our case study. We use the term “variable” loosely to refer to both quantifiable and procedural options in UKF system design. We separate variables that impact the integration of the model equations into the “model variables” class, which can be optimized independently from of the variables that govern the generation of the sigma points. This allows us to remove groups of variants from later optimization tests, thus accelerating the process. The classes of variables and the values we allowed them to take in our optimization are summarized in Table 2, with the benchmark variable values in boldface type.

**Table 2.** UKF system variable classes and values used in case study.

| Abbrev. | Variable | Values |
| --- | --- | --- |
| **Fixed variables** |  |  |
| $m$ | Number of observables | 2 |
| $t_{obs}$ | Observation time interval | 2 s |
| $t_{fh}$ | Forecast horizon time | 10 s |
| **Model variables** |  |  |
| $F$ | Function approximation | **tanh**, LP7, LP3 |
| $M_{int}$ | Integration method | **RK4**, tpz., Eul. |
| $t_{int}$ | Integration time step | **0.25**, 0.5, 1, 2 s |
| $p$ | Numerical precision | **double (d)**, single (*s*) |
| **UKF structure variables** |  |  |
| $N_{sig}$ | Number of sigma points | **2n (24)**, $n + 1$ (13) |
| $M_{sr}$ | Square root method | **Schur**, Chol., SR-UKF |
| - | Matrix inversion | $2 \times 2$ closed form |
| - | Sigma point regeneration | Off |
| - | Sigma point weights | $W_i = 1/N_{sig}$ for all $i$ |
| - | Approach to constraints on states | Enforced on $\hat{x}$ |

Variables in unshaded rows were varied in the optimization. Variables in shaded rows were held fixed. Benchmark variable values are in boldface type.

#### Fixed variables

Fixed variables are those that are largely predetermined by higher-level system considerations and over which the system designer has little or no control. They therefore remain fixed during the optimization. A given system may impose constraints on different variables compared to another system, so the set of fixed variables listed here should be taken only as an example.

The *number of observables, m*, is the dimension of the vector *y*_*k*_ in Eq (2). In many cases, which or how many observables to use is determined by the system requirements. In other cases, the cost-benefit tradeoff for including an observable should be considered. In general, more observables means more information, which theoretically leads to more accurate state estimation. However, due to the matrix operations involved, the computation time of the UKF grows super-linearly with *m*. A further consideration, which we will not discuss here, is the cost on the subject of additional measurements—for example, additional electrodes placed in brain result in additional damage [1]. Additionally, the observability of the estimated states relative to each observable should also be taken into account; the accuracy of UKF estimation of one or more states may depend more or less heavily upon a particular observable or combination of observables [23]. When two or more observables are highly correlated, at least one could be considered for exclusion if the computation time requirements are tight. For our case study, we used two observables (*m* = 2), *F*_*DR*_ and *F*_*R*_, since the dynamics are poorly reconstructed with only one measured firing rate [24], but simultaneously accessing additional DBB model populations in a live animal without damaging critical brain structures is difficult [25].

The *observation time interval, t*_*obs*_, is the time between successive observations in real time. Although the UKF allows for the value of *t*_*obs*_ to vary, we assume it to be constant. On the one hand, *t*_*obs*_ must be small with respect to the rate of divergence of the sigma points from each other. More specifically, within the space defined by the forward iterates of the sigma points, the flow of the dynamics must be sufficiently linearizable. If the dynamics bifurcate between observations such that the mean of the sigma points is no longer close to the expressed dynamics, the state estimate can fail catastrophically. On the other hand, an entire cycle of the UKF-forecasting scheme must compute between observations; therefore, *t*_*obs*_ must be chosen large enough so that at least the fastest variants can be computed on the target platform. In our case study, we assimilated model-generated observations at the interval *t*_*obs*_ = 2 s.

The *forecast horizon time, t*_*fh*_, is how far into the future beyond the next observation the state forecast is made in computed model time. In other words, for every cycle of the UKF-forecasting scheme, the sigma points will be integrated forward by *t*_*obs*_ for the prediction step and an additional *t*_*fh*_ for the forecast. We set *t*_*fh*_ = 10 s because our application involves anticipating changes in state-of-vigilance with a lead time of 5 to 10 s.

#### Model variables

Model variables are those that impact the fidelity with which model integration approximates the dynamics of the idealized model in Eq (1), which in turn tracks the dynamics of the real system with some finite error. *Function approximations* can replace elementary functions inside the model. For example, the tanh function, which appears frequently in neural mass models and can take a relatively long time to compute in MCUs, may be replaced with a linear-piecewise approximation. Taylor series may also be used to approximate functions. In our case study, in addition to the default tanh function from the MCU software library, we considered two linear-piecewise (LP) replacements for the tanh function: a seven-piece approximation of tanh (LP7) and a family of ten three-piece functions (LP3), in which the slope of each three-piece function was individually tuned for the model state in which it was used. The tanh function was used for the benchmark variant.

Various *integration methods, M*_*int*_, and *integration time steps, t*_*int*_, can be tested to find a balance between fidelity to the model and computational cost. A fourth-order Runge-Kutta (RK4) method, for example, will offer higher fidelity to the model ODEs than the simpler trapezoidal or Euler methods but will take longer to compute. (See pp. 29-33 in [22].) Similarly, a smaller *t*_*int*_ will provide higher fidelity than a larger *t*_*int*_ but with increased computation time. We used the RK4, trapezoidal (tpz.), and Euler (Eul.) integration methods, each for *t*_*int*_ = 0.25, 0.5, 1, and 2 s. The RK4 integrator and *t*_*int*_ = 0.25 s were used for the benchmark variant.

The *numerical precision* used—i.e. single- or double-precision floating point or various fixed-point word lengths—can greatly impact computation time on a processor that lacks a floating-point unit (FPU) or has a word size smaller than the chosen data type. MCUs without an FPU can typically perform floating point computations through software routines, but at an increased time cost. We tested both single-precision (32-bit) and double-precision (64-bit) floating point types. Double-precision was used for the benchmark variant.

#### UKF structure variables

We grouped remaining variables that impact any aspect of the UKF, aside from the model, under UKF structure variables. We focused specifically on the *number of sigma points, N*_*sig*_, and the *matrix square root method, M*_*sr*_, applied to 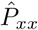.

The standard formulation of the UKF [17] uses a symmetrical set of 2*n* + 1 sigma points, including the mean 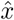, where *n* is the dimension of *x*. The mean can be excluded from the set to reduce the number of sigma points to 2*n*. The spherical-simplex (SS) unscented transform uses a reduced set of *n* + 2 or *n* + 1 sigma points, again depending on whether the mean 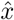 is included [21]. In our case study, we considered both the standard set of 2*n* sigma points and the SS set of *n* + 1 sigma points. We used the standard set for the benchmark.

The formation of sigma points requires a matrix square root *S* of the positive-semidefinite covariance matrix 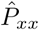, such that 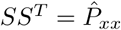. A variety of methods for computing *S*, which in general is not unique, have been applied to the UKF. A common choice for *S* is the lower-triangular factor *L* produced by Cholesky decomposition (Chol.), which has a straightforward closed-form solution. The square root based on the Schur decomposition yields the principal square root, which is symmetrical for a positive semi-definite input matrix and produces sigma points that are generally more evenly distributed in state space than those produced by the Cholesky factor. However, the Schur square root is computationally more intensive than the Cholesky square root. More importantly, the number of operations and therefore time required to compute the Schur square root can vary widely from iteration to iteration within the same code—varying up to 10% in our experience—depending on the condition of the input matrix. This can pose serious problems in a real-time system, and the range of computation times should be checked against the time safety margin to ensure computation deadlines are consistently met. Alternate matrix square root methods for UKFs, not discussed here, are considered in [26] and [27], including iterative methods and methods based on singular-value decomposition.

The square-root UKF (SR-UKF) [19] is theoretically equivalent to employing the Cholesky square root, but it avoids repeatedly computing the Cholesky decomposition of 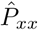 every cycle by tracking the Cholesky factor *L* instead. 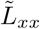 and 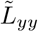, the Cholesky square roots of 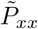 and 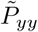, respectively, are obtained through *QR* decomposition followed by a rank-one Cholesky update, if 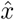 is included as a sigma point. 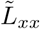 is used to form the sigma points, and 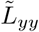 and its transpose are used in back-substitutions to solve for the Kalman gain *K*, which obviates taking the inverse of 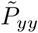. The corrected Cholesky factor 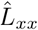 is found through repeated rank-one Cholesky updates of 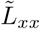. The SR-UKF may provide reduced computation time and better numerical stability compared to the traditional Cholesky method, although the output values of 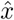 may differ slightly due to the alternative computational approach. We tested the Schur and Cholesky square root methods along with a slightly modified version of the SR-UKF, in which we inverted 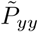 directly rather than using repeated back-substitution to compute *K*, since the dimension of 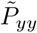 was only 2 in our system. We used the Schur square root for the benchmark case.

A number of other UKF structure variables that we held fixed in our optimization deserve mention. Different strategies for handling the inversion of 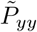 in Eq (9) will have consequences for both computation time and numerical stability, particularly when *m* is large. Because the dimension of our observed state is small (*m* = 2), we used an explicit solution for a 2 × 2 matrix inversion. The sigma points may be regenerated after the prediction step and before transformation into the observation space [22]. The impact of this step on state estimation accuracy may depend on the degree of nonlinearity in the observation function Eq (2). We bypassed sigma point regeneration after the prediction step. The sigma point weights *W*_*i*_ can be adjusted not only to include or exclude 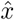 from among the sigma points but also to incorporate prior knowledge of the higher-order moments of the state distribution [17, 19]. Methods for tuning the weights are described in [28–32]. We weighted the sigma points by the reciprocal of their total number. Finally, the noise covariance matrices *Q* and *R* may also undergo tuning, as addressed in [28, 31, 33]. We held *Q* and *R* constant and inflated the element in *Q* corresponding to the uncertainty of the unknown input *δ*.

In general, it is useful to check if generated sigma points fall within the allowed state space of the model dynamics and, presumably, the real system. For example, real neuronal systems do not have negative firing rates. Methods have been developed for dealing with constraints on state values in both linear and nonlinear Kalman filters [22, 34–38]. Although the states in the DBB model are physically meaningful only for non-negative values, the state estimation accuracy of our system was not found to be significantly impacted by small excursions of the sigma points into negative territory.

From the model equation side, this can be understood because steady-state firing rates in Eq (14) are formulated to be positive definite despite potential negative neurotransmitter concentrations, and the steady-state neurotransmitter concentrations in Eq (17) produce positive outputs from positive firing rate inputs. Therefore the only accommodation we made for the non-negative state constraints was to enforce them on 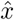.

### Computation time test

In the computation time test, we identify whether each variant of the UKF prediction-correction-forecast scheme can be computed in real-time. For of each of the 432 combinations of model and UKF structure variables represented in the unshaded rows of Table 2, we measured the computation time *t*_*comp*_ for one UKF cycle on the target MCU. In Fig 3 we depict the sequential computational tasks comprising *t*_*comp*_, beginning with the moment the observed data *y*_*k*_ become available: the observed data are used to correct the predicted mean and covariance from the previous cycle; the new set of sigma points are generated; the sigma points are propagated forward by *t*_*obs*_ + *t*_*fh*_ in model time; and the predicted mean and covariance are computed for the next cycle. Note that the computation time test assumes that the specific target processor and values of *t*_*obs*_ and *t*_*fh*_ are fixed.

**Fig 3.**
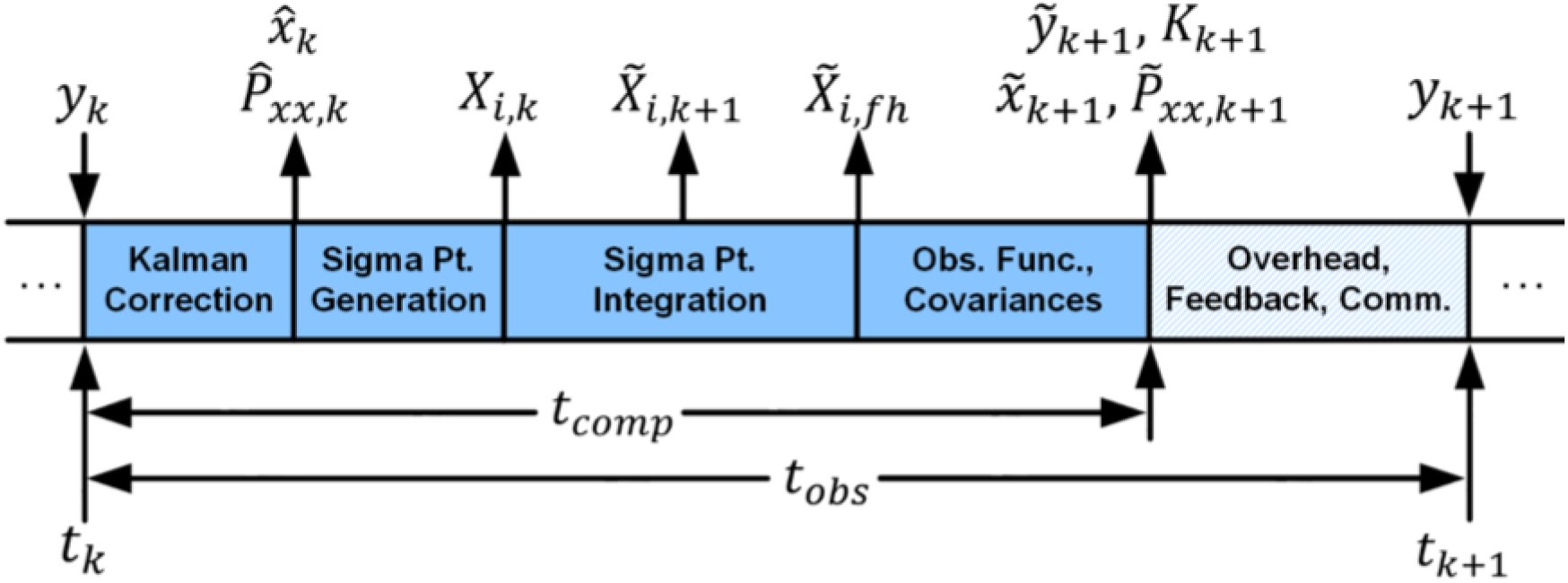
UKF system computational tasks in time relative to *t*_*obs*_. One cycle of the UKF must compute in the time between successive observations, *t*_*obs*_, while also allowing for overhead tasks, such as feedback to the biological system and communication with a base station. The computation time *t*_*comp*_ refers to the time required to compute the UKF matrix operations and generate and integrate the sigma points. Block widths are not to scale.

For each variant, *t*_*comp*_ was obtained by computing two cycles of the UKF prediction-correction-forecast scheme on the target MCU and capturing the number of clock periods required to compute the *second* UKF cycle. Because *t*_*comp*_ is relatively invariant to the location of 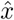 in state space, the timing test only requires computation of two cycles of the UKF, rather than hours of data in model time. This enables collection of *t*_*comp*_ data directly from the MCU, which eliminates the need for clock-cycle-accurate MCU emulation software for the PC. The total time required to obtain *t*_*comp*_ for all 432 variants was 17 min. The C codes used to obtain the values of *t*_*comp*_ on the Cortex-M3 were translated from MATLAB using MATLAB Coder.

The second and subsequent cycles of the UKF compute more slowly than the first cycle. This difference is due in part to the increased time spent in the square root operation on the non-diagonal matrix 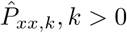, compared to the faster square root performed on the diagonal initial-condition matrix 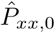 in the first cycle. It should be noted that some variability in *t*_*comp*_ is expected for matrix functions containing operations that are conditional on the qualities of the input matrix, like the Schur square root and QR decomposition. For our case study, one value of *t*_*comp*_ is sufficient, since the variation in time to compute these functions is very small relative to our time safety margin. For a different system, further probing of the range of computation times may be necessary.

In the optimization, measured values of *t*_*comp*_ for each variant are compared to a maximum threshold value *θ*_*t*_ *< t*_*obs*_. Variants with *t*_*comp*_ *> θ*_*t*_ fail the computation time test. The threshold *θ*_*t*_ is constrained from above by not only *t*_*obs*_ but also by the time required for overhead tasks (*t*_*oh*_) and a safety margin (*t*_*marg*_). Thus, *θ*_*t*_ ≤ *t*_*obs*_ − *t*_*oh*_ − *t*_*marg*_. Cases may arise in which feedback to the subject resulting from the forecast must be initiated before the end of the *t*_*obs*_ interval. In such cases, it may be necessary to also measure the computation time from the moment of observation to the moment when the forecast is known (*y*_*k*_ to 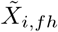 in Fig 3). For this study, we chose *θ*_*t*_ = 1.5 s, which allows 0.5 s for *t*_*oh*_ and *t*_*marg*_, combined, and also satisfies our feedback latency requirement.

It should be emphasized that *t*_*comp*_ ≤ *θ*_*t*_ is merely a necessary condition. In general, a smaller computation time is preferable for at least three reasons: smaller computation time implies 1) lower power and 2) lower feedback latency, and 3) a larger margin between *t*_*comp*_ and *θ*_*t*_ allows for greater flexibility in modifying the system design at a later time without violating timing constraints.

The impact of the model and UKF variables on *t*_*comp*_ will be more or less significant depending on where the variant sits in the optimization variable space. The computation time may be approximated as

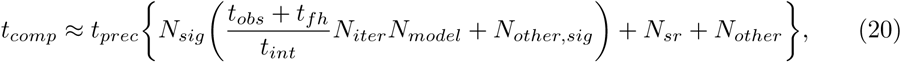

where *t*_*prec*_ is the average time for an operation of a given precision, *N*_*sig*_ is the number of sigma points used in a given UKF, *N*_*iter*_ is the number of model equation iterations used in a given integration method, *N*_*model*_ is the number of operations in the model equations, *N*_*other,sig*_ is the number of other operations repeated for each sigma point, *N*_*sr*_ is the number of operations in the matrix square root, and *N*_*other*_ is the number of operations not captured elsewhere. For example, a change in *t*_*comp*_ due to the matrix square root variant will only manifest when the *N*_*sig*_ term is on the same order as *N*_*sr*_ or smaller.

The high-level trends implied by Eq 20 are reflected in Fig 4A, where we show computation time test results for the benchmark UKF system and all variants that differ from the benchmark by only one variable. The change from double to single precision reduces *t*_*comp*_ by a factor of about two. The dominance of the *N*_*model*_ term in this region of optimization variable space is evidenced by 1) the halving of *t*_*comp*_ for each doubling of *t*_*int*_; 2) the proportionality of *t*_*comp*_ to *N*_*iter*_, where *N*_*iter,RK*4_ = 4, *N*_*iter,tpz*_ = 2, and *N*_*iter,Eul*_ = 1; and 3) the proportionality of *t*_*comp*_ to *N*_*sig*_, where *N*_*sig*_ = 24 for the benchmark and *N*_*sig*_ = 13 for the SS variant. The significant time cost of evaluating the tanh function in the model equations is shown by the roughly 60% reduction in *t*_*comp*_ when the linear piecewise approximations LP7 and LP3 are used.

**Fig 4.**
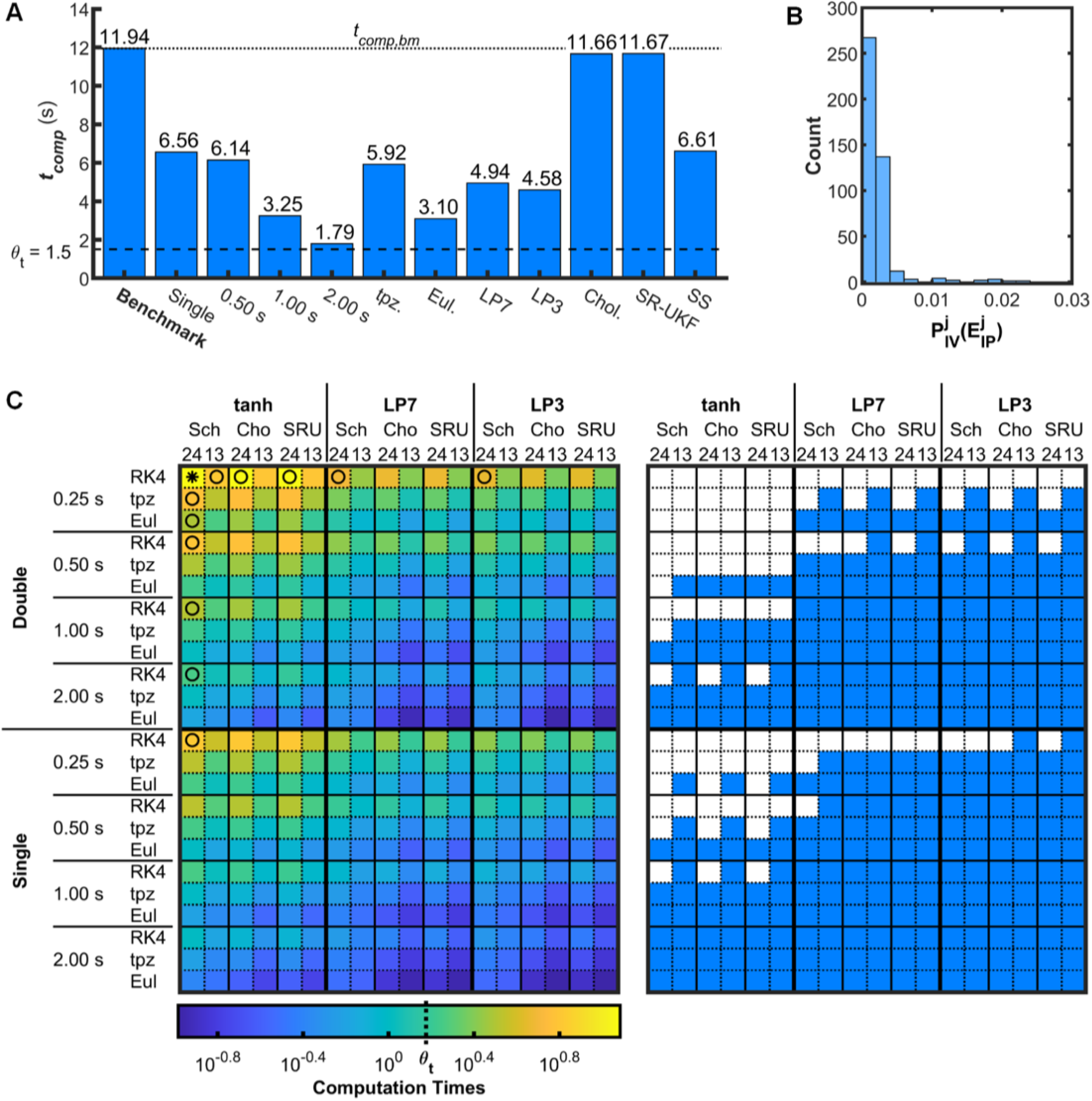
Computation time test results and platform validation. (A) Values of *t*_*comp*_ for the benchmark variant (*t*_*comp,bm*_, dotted line) and all variants that are one variable step away from the benchmark. Labels indicate the differing variables. (B) Histogram of 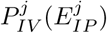 values for platform validation. (C) *t*_*comp*_ is shown for all variants. (Left) *t*_*comp*_ color coded. The benchmark variant (*) and other variants (◦) for which *t*_*comp*_ is plotted in (A) are marked. (Right) Same data binary thresholded for *t*_*comp*_ ≤ *θ*_*t*_. Abbreviations: seven-piece linear (LP7) and three-piece linear (LP3) approximations of tanh; 4th-order Runge-Kutta (RK4), trapezoidal (tpz), and Euler (Eul) integration methods; Schur (Sch) and Cholesky (Cho) square root methods; square-root UKF (SRU); standard (24) and spherical-simplex (13) sigma point methods.

Although none of the variants in Fig 4A compute in less than *θ*_*t*_ = 1.5 s, many variants—about three quarters of those tested—passed under the threshold, with the fastest variant computing in 0.105 s, compared to 11.94 s for the benchmark variant. The values of *t*_*comp*_ for all variants are represented on the left side of Fig 4C, with the passing variants having *t*_*comp*_ *< θ*_*t*_ shaded in the grid on the right. For variants in which the *N*_*sig*_ term has been substantially decreased, the impact of the faster square root methods on *t*_*comp*_ become apparent. For example, compare the similarity of *t*_*comp*_ between the Schur and Cholesky methods in the top row of Fig 4C (left) to the noticeable difference between the Schur and Cholesky methods in the bottom row.

### Forecast test

In the forecast test we identify whether the computed forecast trajectories—uncorrected by observed data—for a given combination of model variables follow benchmark system trajectories with sufficient fidelity.

The quantitative criterion used in the forecast test is rooted in the probabilistic description of system state adopted in KFs. The distribution of state is centered at 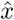 with “width” 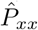. Predictions and forecasts are made by evolving this distribution through the model equations. As 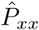 evolves forward in time, it gradually expands due to the process noise *Q*, which encapsulates the unmodeled dynamics. Our objective is to identify alternate methods of integrating the model equations—i.e. using simpler integrators, larger time steps, faster precision, and model approximations—that maintain sufficient fidelity to the benchmark integration scheme relative to *Q*. We define fidelity through a measure of how far trajectories integrated with these variant methods diverge from trajectories computed with the benchmark integrator over the anticipated forecast time *t*_*fh*_. The variant and benchmark trajectories are computed with zero process noise; thus the divergence between them is due to the computational simplifications made in the variants. To assess this variant-to-benchmark forecast divergence error 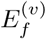 as it evolves over time, we compare it with the process noise *Q*, which we obtain by computing the same divergence error metric between benchmark-integrated trajectories without noise and “true” noisy data 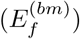.

The data for both the forecast and state reconstruction tests were computed in MATLAB running on a desktop PC, rather than on the target MCU, due to the large volume of data. Because the two platforms use different programming languages, compilers, processors, and instruction sets, an inter-platform error 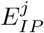 is expected between the output from the Cortex-M3 for the *j*^th^ UKF variant and output from MATLAB running on the PC for the same variant. In order for the proxy PC/MATLAB data for the *j*^th^ UKF variant to be valid in assessing its performance on the Cortex-M3 relative to other variants, 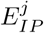 should be small compared to the inter-variant errors 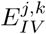, which are the differences between the output of the *j*^th^ and *k*^th^ UKF variants, *k* ≠ *j*, computed on the Cortex-M3.

We expect that 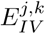 for some pairs of variants will be close to zero. However, the goal of the optimization stages is to identify and remove outlying variants that fail in *significant* ways. Therefore 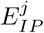 for the *j*^th^ variant need only be smaller than the median of its 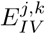 values. Furthermore, if methods *j* and *k* have near-zero 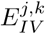 on the target platform, their numerical performance is equivalent for our purposes and they need not be distinguished from each other.

To validate the PC/MATLAB as a substitute computational platform for the Cortex-M3, we computed two cycles of the UKF—the first with a diagonal initial-condition 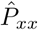 matrix, and the subsequent iteration with a non-diagonal 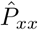 matrix—for every tested variant both on the Cortex-M3 (target) and on the PC with MATLAB (proxy). We retained the state estimation outputs 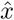 from the second cycle and computed the inter-platform error

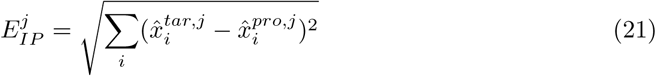

and inter-variant error

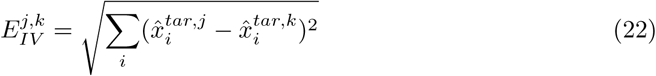

for all *j* and all *k* ≠ *j*, where 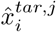 and 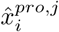 are the *i*^th^ entry of the 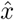 output vector for the *j*^th^ variant computed on the target platform and proxy platform, respectively.

For variant *j*, we define the empirical cumulative distribution function (ECDF) of its inter-variant error 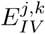 over all *k* ≠ *j* as 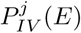, which is bounded on the interval [0, 1]. We require that a variant’s inter-platform error be less than the median of its inter-variant error, or 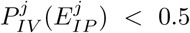. This condition was met for all of the tested variants. In fact, for all variants 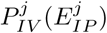 was less than 0.025, which indicates that MATLAB running on the PC performs sufficiently well as a proxy for UKF computations on the Cortex-M3—in both double and single precision—for this case study. A histogram of the 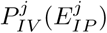 values for all *j* is shown in Fig 4B.

For the forecast test, a large set of synthetic “true” noisy data (5000 min) was generated in MATLAB using the benchmark model variables: the tanh function, an RK4 integrator with *t*_*int*_ = 0.25 s, and double precision. This true data included the random input *δ* and the neurotransmitter noise. In this simulated context, *δ* and the neurotransmitter noise comprise the process noise. A set of 10 000 initial conditions were randomly selected from the true data, and each starting point was integrated forward for *t*_*obs*_ + *t*_*fh*_ = 12 s without process noise by the benchmark integrator and the integrator corresponding to each combination of model variables that passed the computation time test.

For each of the 10 000 test windows, the normalized forecast error 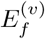 for each variant *v* was found between the end point of the 12-s-long variant-integrated trajectory 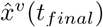 and the end point of the 12-s-long benchmark-integrated trajectory 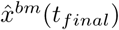:

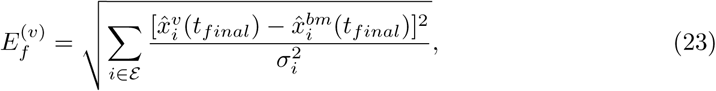

where *σ*_*i*_ is the standard deviation for model state *i*, and ℰ is the set of model states critical to the application (here, ℰ = {DR, VLPO, R}). This yielded a distribution of 10 000 values of 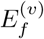 for each tested model variant. The normalized forecast error between the noiseless benchmark-integrated trajectories and the true noisy data 12 s after each initial condition, 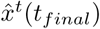, is similarly defined:

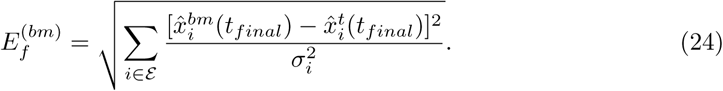

The distribution of 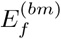 over the 10 000 test windows was taken as the process noise *Q*, against which the variants were assessed.

Each combination of the *model* variables in Table 2 corresponds to a set of six variants of the UKF *structure* variables: two options for the number of sigma points (*N*_*sig*_) multiplied by three options for the square root method (*M*_*sr*_). The forecast test was run for the combinations of model variables (*F, M*_*int*_, *t*_*int*_, and *p*) that had at least one out of the six corresponding UKF structure variants pass the computation time test.

In order to assess the fidelity of a tested variant to the benchmark, we compare the ECDF of 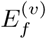, defined as *P*_*f,v*_(*E*_*f*_), for each variant under test to the ECDF of 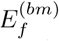, defined as *P*_*f,bm*_(*E*_*f*_); in other words, we compare the “width” of the spread due to the simplified variant to the “width” of the spread due to *Q*, both at the forecast time horizon. In the context of the UKF, the state distribution iterated forward in time produces variances that are ideally accommodated by the UKF correction step. However, prediction-correction schemes perform poorly with large-outlier predictions that place the estimated state, for example, across trajectory bifurcations. Therefore in order to focus the forecast test metric on these problematic large-outlier cases, we compare the ECDFs at the 95^th^ percentile using a right-side tail comparison metric *T*_*f*_ :

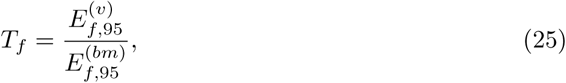

where 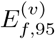 and 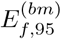 are the 95^th^ percentile values of *P*_*f,v*_ and *P*_*f,bm*_, respectively. We set the maximum threshold of the tail metric *T*_*f*_ for the forecast test, *θ*_*f*_, to 1. Model variants with *T*_*f*_ ≤ *θ*_*f*_ = 1 are considered to have forecasting fidelity to the benchmark integrator comparable to or better than the fidelity of the noiseless benchmark integrator to the true noisy system. Variants with *T*_*f*_ *> θ*_*f*_ fail the forecast test. Note that the failure of a single *model* variant eliminates up to six UKF *structure* variants from the following test.

Sixty of the seventy-two model variants passed the timing test and were assessed in the forecast test. The ECDFs *P*_*f,v*_ for selected model variants are shown in Fig 5A along with *P*_*f,bm*_. Box charts of the same *E*_*f*_ data are shown in Fig 5B along with the corresponding values of *T*_*f*_. In Fig 5C, values of *T*_*f*_ for all variants are represented in the left half, and variants that passed under the threshold *θ*_*f*_ = 1 are shaded in blue in the right half. The red box highlights the variable space corresponding to the data plotted in Figs 5A and 5B.

**Fig 5.**
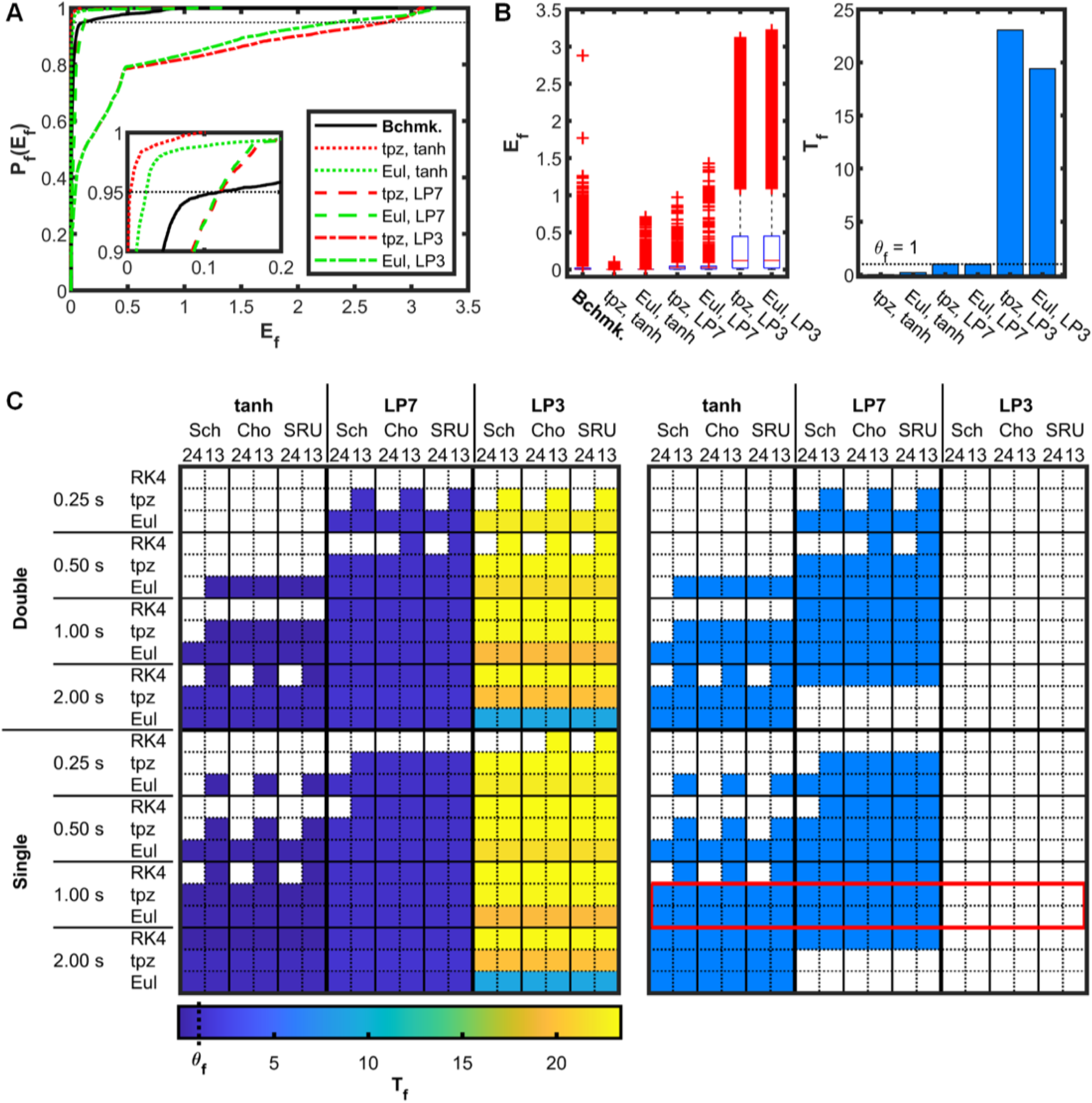
Forecast test results. (A) The benchmark variant ECDF *P*_*f,bm*_ is plotted with *P*_*f,v*_ for the model variants that use single precision, *t*_*int*_ = 1 s, and trapezoidal or Euler integration. The neighborhood where the benchmark and four of the other ECDFs enter the 95^th^ percentile, marked by the horizontal dotted line, is enlarged in the inset. (B) Box charts of *E*_*f*_ and values of *T*_*f*_ for the same variants. (C) *T*_*f*_ is shown for all variants that passed the timing test. (Left) *T*_*f*_ color coded. (Right) Binary thresholded for *T*_*f*_ ≤ *θ*_*f*_. The red box highlights the model variants for which data are presented in (A) and (B).

Replacing the tanh function by the LP3 caused the forecast to fail with a magnitude of error generally much larger than the magnitude of the process noise. Integration method and integration time step played smaller roles in decreasing fidelity of the variants to the benchmark. Use of single precision in place of double precision had a negligible impact on the fidelity.

The variable space in Figs 4C and 5C is laid out such that *t*_*comp*_ generally decreases from top to bottom and from left to right. The computation time test (Fig 4C) eliminated more time-intensive variants from the upper and left regions of the double and single portions of the variable space. The forecast test (Fig 5C) eliminated less time intensive variants from the lower and right regions of the double and single portions of the variable space.

Some unexpected results highlight a point of caution. In Fig 5C (left) the apparent improvement in results for the Euler integration of the LP3 variants at *t*_*int*_ = 2 s over the RK4 and trapezoidal methods is misleading. Analysis revealed oscillations due to instability for those variants at certain locations in state space. The oscillations pushed the state forecast away from and then back toward the initial-condition region of state space every two integration time steps. Thus the even number of integration time steps over a short total integration time resulted in the error unexpectedly decreasing compared to stable integrations involving the RK4 and trapezoidal methods, under those conditions. This example underscores the importance of data visualization across the entire variable space in order to identify potentially misleading results.

The forecast test eliminated sets of six variants sharing the same model variables. In the state reconstruction test, the impact of the UKF structure variables on state reconstruction is examined.

### State reconstruction test

In the state reconstruction test, we identify if the mean state estimate 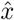 produced by a given UKF variant tracks the benchmark UKF 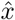 with sufficient fidelity. Here “sufficient fidelity” is defined by the distribution of error between the benchmark UKF state estimate and the true state, against which we assess the error between the output of each variant and the benchmark UKF.

We generated 1000 one-hour-long segments of “true” noisy data using the benchmark model integrator and including the random input *δ* and the neurotransmitter noise, as with the true data for the divergence test. Synthetic noisy observation data were generated for the observed variables (*F*_*DR*_ and *F*_*R*_) by adding Gaussian noise to the true data. Negative observation values were corrected to zero. We then seeded the benchmark UKF and each surviving UKF variant with the true initial conditions and used them to reconstruct state estimates using the noisy observation data for the 1000 segments. These tests were computed in MATLAB.

In a manner similar to the forecast test, the error between each of the tested variants and the benchmark UKF was assessed relative to the error between the benchmark UKF and the true data. For each tested variant, we computed the normalized mean-squared error between the state estimates reconstructed by the variant UKF and the benchmark UKF,

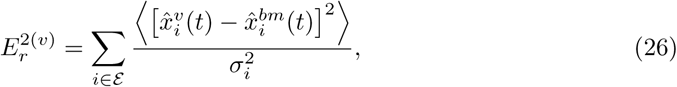

for each of the 1000 test windows, with the mean taken over the hour-long window. This yielded a distribution of error for each variant. We used an analogous formula to compute the normalized mean-squared error between the benchmark UKF data and the true data, 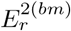.

Whereas in the forecast test the process noise *Q* provides the leeway for simplifying the model integration, the constraining influence of the UKF correction step provides the leeway for simplifying the UKF structure in the state reconstruction test. Once every *t*_*obs*_, the estimated distribution of state is pulled toward the observed data using the Kalman gain in Eq (11). As long as the predicted cluster of sigma points approximately encompasses the true mean, the correction step will compensate for the simplifications in both the UKF structure and the model integration. However, if the forward iterated sigma points diverge from the true state in one or more dimensions, the impact of the correction step may be insufficient to prevent the entire sigma point set from drifting into catastrophic error. Therefore we again focus the test metric on these problematic large-outlier cases by comparing the reconstruction error ECDFs at the 95^th^ percentile:

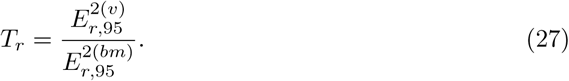

Variants with *T*_*r*_ *> θ*_*r*_ = 1 fail the state reconstruction test.

We define the ECDF of 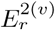 as 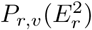, which is plotted in Fig 6A for selected variants against the ECDF of 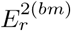, which we define as 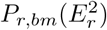. The corresponding box charts and *T*_*r*_ values are shown in Fig 6B. Values of *T*_*r*_ for all variants are represented in the left side of Fig 6C, while passing variants are shaded in blue on the right side. The red box in Fig 6C (right) highlights the variable space corresponding to the data plotted in Figs 6A and 6B.

**Fig 6.**
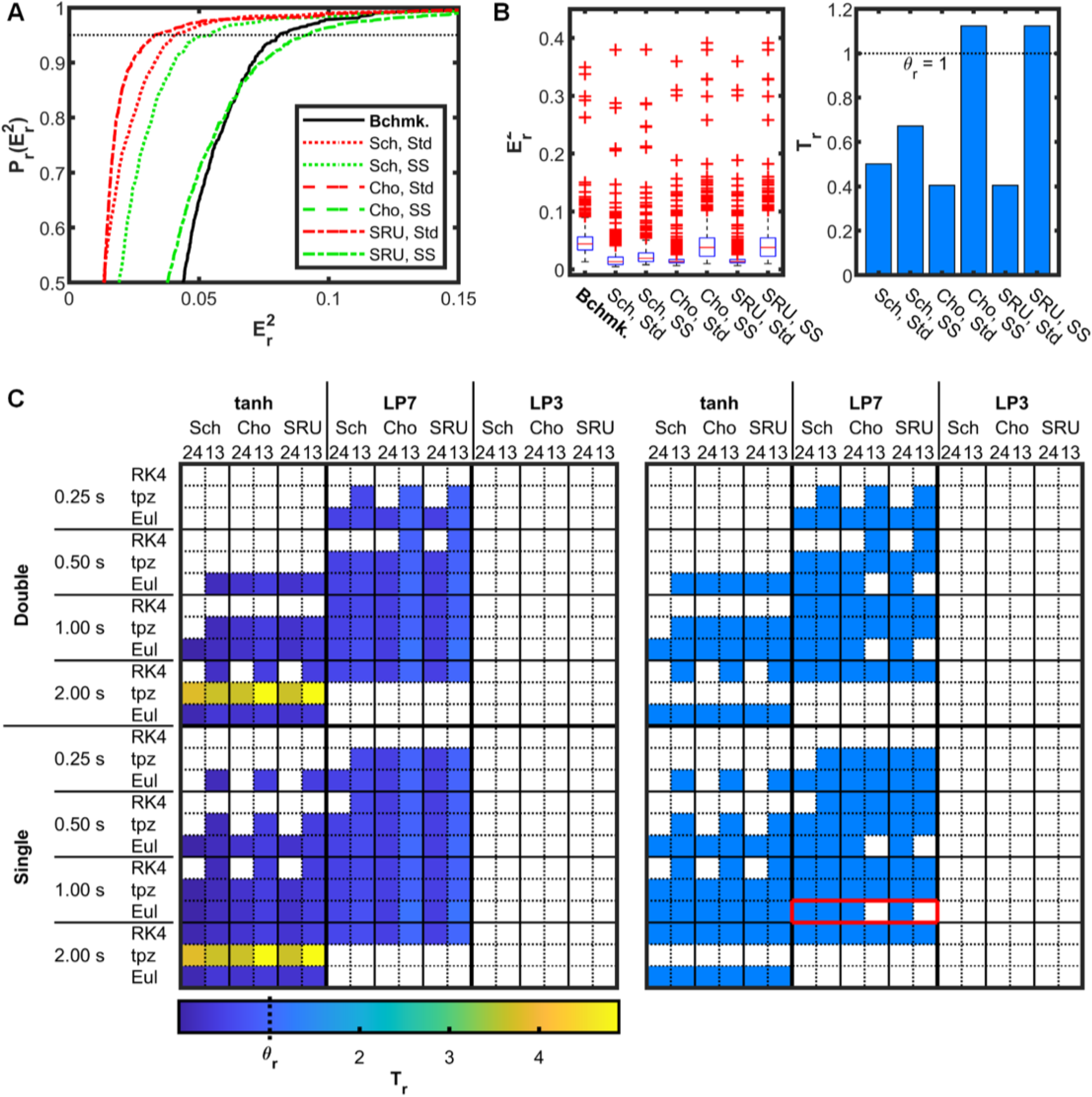
State reconstruction test results. (A) The benchmark variant ECDF *P*_*r,bm*_ is plotted with *P*_*r,v*_ for the model variants that use single precision, *t*_*int*_ = 1 s, Euler integration, and the LP7 approximation. The horizontal dotted line marks the 95^th^ percentile. Note that the ECDFs for the SR-UKF variants overlap those of the Cholesky square root variants. (B) Box charts of 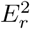 and values of *T*_*r*_ for the same variants. (C) *T*_*r*_ is shown for all variants that passed the forecast test. (Left) *T*_*r*_ color coded. (Right) Binary thresholded for *T*_*r*_ ≤ *θ*_*r*_. The red box highlights the variants for which data are presented in (A) and (B).

In Fig 6 the similarity in performance between the Cholesky (Cho) and SR-UKF (SRU) square root methods is apparent. In fact the SR-UKF ECDFs entirely overlap those of the Cholesky square root method. It should not be assumed, however, that this will be the case for all systems. Changes in precision, state dimension, and the underlying model could each potentially draw out differences in the numerical approaches between these two methods.

For all square root methods, the standard set of 24 sigma points (Std) tends to outperform the reduced set of 13 sigma points (SS). However, within families of six variants having the same model variables, when 24 sigma points are used the Cholesky square root and SR-UKF methods tend to outperform the Schur square root method, and when 13 sigma points are used, the opposite is true.

We hypothesize that these trends are due to the coupling of nonlinearities in the model with the slight differences in sigma point arrangement produced by the different square root and sigma point generation methods. For a given distribution with mean 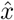 and covariance 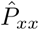, all sigma point generation methods and matrix square root methods produce sigma points with the same mean and covariance. Furthermore, for all methods, sigma points lie on the same isoprobability. However, the “coverage” of state space differs among methods. For example, because the Cholesky square root is lower triangular, it systematically produces a set of sigma points with a larger span—relative to the variance—in states with lower dimension index than states with higher dimension index, whereas the symmetrical Schur square root tends to produce sigma points that are similarly distributed among all states. Also, regardless of the matrix square root method, the standard method produces sigma points that are arranged in pairs opposite from each other in state space, with the mean 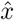 in between. In contrast, the spherical-simplex method produces a sigma point cloud lacking this form of symmetry.

The trapezoidal integration method for *t*_*int*_ = 2 s fails in the state reconstruction tests. In the DBB model, trapezoidal integration with a large *t*_*int*_ can lead to an unusual situation in which *F*_*R*_ increases to its typical “on” firing rate while the acetylcholine emitted by the REM-active sub-population (*C*_*AR*_) remains low. This appears to be due to negative feed back that occurs under certain conditions in the *C*_*AR*_ equation from the first to the second evaluation of the equation in the trapezoidal method. This unexpected result highlights the importance of sufficiently sampling the model state space to draw out such anomalies.

The insensitivity to double versus single precision for this model and selected UKF variable space is evident in the similarity between the top and bottom halves of Fig 6C. Precision may become relevant, however, for stiffer model equations or a higher-dimension of the matrix 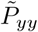, which undergoes inversion.

### Cost function

A multi-objective cost function combining the results of the optimization stages was applied to the variants that passed all three tests. The cost function is a sum of three weighted terms, one each for computation time, forecast error, and state reconstruction error:

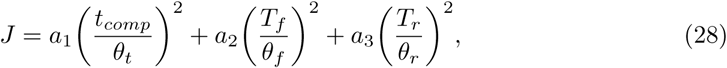

where *a*_1_, *a*_2_, and *a*_3_ are adjustable weights (set to *a*_1_ = *a*_2_ = *a*_3_ = 1 for this case study). Since Eq (28) is a weighted sum of individual cost functions to be minimized, the variant that minimizes Eq (28) is guaranteed to be Pareto-optimal. Squared terms are used to flatten the cost function near the minimum.

In most MCU-based applications, power is roughly proportional to *t*_*comp*_. However, if FPGA fabric is used to realize part of the system design and power data are available for the tested variants, a fourth term my be added to Eq 28 that incorporates computation power *P*_*comp*_ and a corresponding threshold *θ*_*P*_. Power data can be used to eliminate variants in a separate test similar to the timing test. *P*_*comp*_ was not separately considered in our MCU-based case study.

The ten variants that yielded the lowest values of the cost function Eq (28) are listed in Table 3, and cost function values for all UKF system variants that passed the state reconstruction test are indicated in Fig 7. The optimal variant with the lowest cost function value (*p* = *s, t*_*int*_ = 1 s, *M*_*int*_ = Euler, *F* = tanh, *M*_*sr*_ = Schur, *N*_*sig*_ = 13) computes in 0.439 s, which is 27 times faster than the benchmark variant (see Fig 4A).

**Table 3.** Variants with the lowest cost function values.

| | CF | $t_{comp}$ (s) | $T_f$ | $T_r$ | $p$ | $t_{int}$ (s) | $M_{int}$ | $F$ | $M_{sr}$ | $N_{sig}$ |
| --- | --- | --- | --- | --- | --- | --- | --- | --- | --- | --- |
| 1 | 0.150 | 0.439 | 0.200 | 0.157 | $s$ | 1.0 | Eul. | $\tanh$ | Schur | 13 |
| 2 | 0.186 | 0.455 | 0.200 | 0.232 | $s$ | 1.0 | Eul. | $\tanh$ | SR-UKF | 24 |
| 3 | 0.187 | 0.458 | 0.200 | 0.232 | $s$ | 1.0 | Eul. | $\tanh$ | Chol. | 24 |
| 4 | 0.197 | 0.637 | 0.033 | 0.126 | $s$ | 1.0 | tpz. | $\tanh$ | Schur | 13 |
| 5 | 0.206 | 0.612 | 0.200 | 0.012 | $s$ | 1.0 | Eul. | $\tanh$ | Schur | 24 |
| 6 | 0.215 | 0.649 | 0.100 | 0.134 | $s$ | 0.5 | Eul. | $\tanh$ | Schur | 13 |
| 7 | 0.229 | 0.647 | 0.041 | 0.203 | $s$ | 2.0 | RK4 | $\tanh$ | Schur | 13 |
| 8 | 0.238 | 0.281 | 0.200 | 0.404 | $s$ | 1.0 | Eul. | $\tanh$ | SR-UKF | 13 |
| 9 | 0.239 | 0.283 | 0.200 | 0.404 | $s$ | 1.0 | Eul. | $\tanh$ | Chol. | 13 |
| 10 | 0.273 | 0.490 | 0.033 | 0.407 | $s$ | 1.0 | tpz. | $\tanh$ | SR-UKF | 13 |

**Fig 7.**
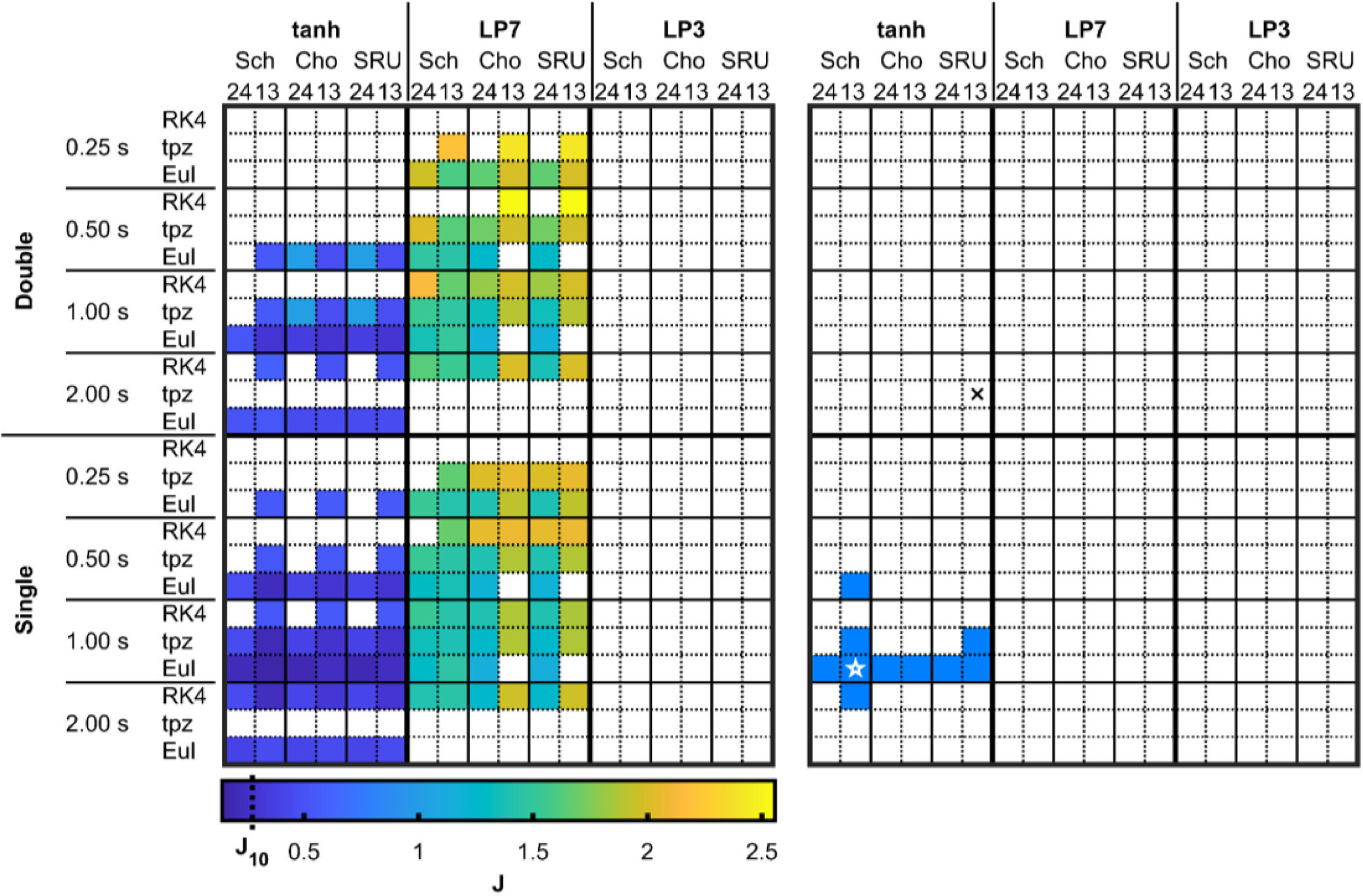
Cost function values. (Left) Values of the cost function *J* are represented in color for variants that passed all three optimization tests. Variants that failed any of the three tests appear as white. *J*_10_ denotes the tenth-lowest cost function value. (Right) The ten variants with the lowest cost function values are shaded in blue. Reconstructed data for the optimal variant (⋆) and a variant that failed the reconstruction test (×) are shown in Fig 8.

Because the cost function contains multiple competing terms that balance computation speed with fidelity to the benchmark UKF, a variety of variable combinations yielded cost function values less than 1. The ten best variants in Table 3 span all three integration methods, all three matrix square root methods, both sigma point generation methods, and three of the four values of *t*_*int*_. This indicates that a balance between speed and fidelity can be obtained through multiple different approaches.

Six of the ten best variants share the same model variables (*p* = *s, t*_*int*_ = 1 s, *M*_*int*_ = Euler, *F* = tanh) and thus also share the same value of *T*_*f*_. Four of the ten are adjacent pairs of Cholesky and SR-UKF variants. In both pairs, the SR-UKF computes in slightly less time than the corresponding Cholesky variant.

Absent from Table 3 are variants using double precision or the LP7 function. The lowest cost function value for a variant using double precision was 0.284, which ranks 14^th^ among all variants. All of the surviving tanh variants had lower cost function values than the LP7 variant with the lowest cost function value (*J* = 1.150).

In Fig 8, we compare UKF state reconstruction for the optimal UKF and a selected UKF variant that passed the forecast test but failed the state reconstruction test. Data are shown from the window that yielded the 95^th^ percentile 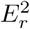 value for the failing variation. The failing variant catastrophically diverges from the true data at the end of the segment and only loosely tracks the downstrokes of *F*_*R*_, whereas the optimal variant maintains acceptable fidelity over the entire hour and closely tracks *F*_*R*_.

**Fig 8.**
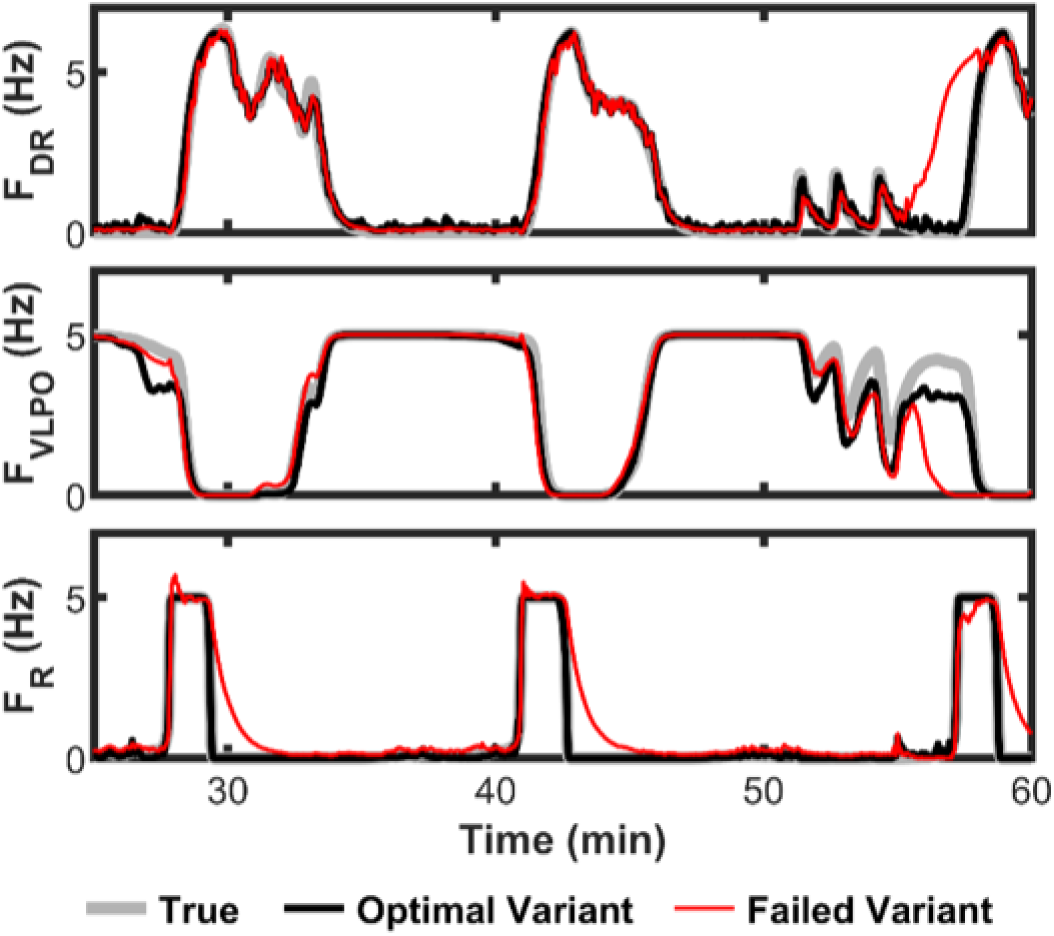
UKF-reconstructed data. True data and UKF reconstructions of the three states used for the error metrics are shown for the optimal variant and a selected variant that failed the state reconstruction test. The data are excerpted from the state reconstruction test window corresponding to 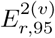 for the failing variation, which uses double precision, *t*_*int*_ = 2 s, trapezoidal integration, the tanh function, the SR-UKF, and spheical-simplex (13) sigma points.

### Optimization computation time

The total computation time for the optimization was 139 h. The values of *t*_*comp*_ were generated from tests run on the target MCU, which took 17 min. The proxy PC was used to generate the forecast and reconstruction test data, which took 8 min and 138 h to generate, respectively. We summarize these times along with the number of variants assessed in each test in Table 4.

**Table 4.** Optimization stage compute times.

| Test | Variants tested | Time | Platform | Variants eliminated |
| --- | --- | --- | --- | --- |
| 1. Computation time | 432 | 17 min | MCU | 105 |
| 2. Forecast | 327 | 8 min | PC | 152 |
| 3. State reconstruction | 175 | 138 h | PC | 20 |

The order chosen for the optimization stages minimizes the time required to compute the results for multiple reasons. The results from the computation time test can be obtained quickly by integrating one or two cycles of the UKF prediction-correction-forecast scheme, making this data relatively inexpensive to acquire over the full variable space at the outset. Placing the computation time test first also disqualifies the most time-intensive variants from later stages. Furthermore, the forecast test decouples the model variables from the UKF structure variables, thereby allowing a family of variants—six in our case—to be assessed together as a unit. And for this case study, the state reconstruction test was by far the most time intensive. Thus, placing it after the other two tests minimized the number of variants that passed through it.

The forecast test involves integration of the model equations without correction from the UKF, thus allowing weaknesses in model variable combinations to become apparent. In contrast, the state reconstruction test includes the UKF correction step, which may mask weaknesses in variables associated with model integration. Therefore it is reasonable to assume that inadequate combinations of model variables are more likely to fail during the forecast test with uncorrected integration than during the state reconstruction test. However, combinations of model variables may potentially couple with combinations of UKF variables to highlight weaknesses in both. The assumption underlying sigma-point KFs is that the distribution of state is well defined by a multivariate Gaussian distribution. The UKF structure variables systematically determine sets of sigma points that represent the state distribution more or less effectively. Meanwhile, the model variables—different integrators and function approximations, for example—alter the shape of the forward integrated distribution with respect to the “ideal” state distribution. It is not guaranteed that these two effects—shortcomings in the arrangement of sigma points and simplification of the integrated dynamics—will not systematically align to expose weaknesses of both. This highlights the importance of testing all surviving combinations of model variables with all of the UKF structure variants in the state reconstruction test.

MATLAB code for the optimization method is available on Penn State’s ScholarSphere at https://doi.org/doi:10.26207/bzgp-ph78.

## Conclusion

We have presented a method for designing an embedded UKF system for a computationally constrained target device. The method seeks an optimum balance between fast computation time on the one hand and fidelity of state forecasting and reconstruction relative to a benchmark UKF on the other hand. We have described an optimization space of UKF system variables and a series of tests to assess UKF system performance in computation time, forecasting, and state reconstruction. We applied the optimization method to an embedded UKF case study that tracks the rat sleep-wake regulatory system. The original benchmark UKF in the case study computed a two-second observation cycle in 11.94 s on the target platform. After applying the series of tests, we ranked passing variants using a multi-objective cost function and presented sample reconstructions comparing the optimal UKF to the benchmark UKF. The optimal UKF reduces the computation time on the target platform by a factor of 27 compared to the benchmark UKF, and it maintains a high level of fidelity to the benchmark reconstruction relative to the benchmark error.

We again emphasize the two aspects of the UKF that allow the computations to be simplified while maintaining relative accuracy: 1) the magnitude of the process noise, which provides leeway for reducing the complexity of the forecast integration, and 2) the influence of the Kalman correction step, which constrains the state estimate error in spite of simplifications made to the UKF structure. We also highlight that the optimization process was accelerated by evaluating the model variables in isolation from the UKF structure variables during the forecast test and ordering the tests to eliminate as many variants as possible from the time intensive state reconstruction test.

Embedded systems are *ad hoc* by nature, and the presented framework should be modified or expanded as needed, for example, by the addition of fixed point types in the precision variable. However, the underlying themes developed in our method should remain applicable across a broad range of applications.

Future work includes deploying an embedded UKF system to predict sleep-state transitions for *in vivo* epilepsy experiments in rat models. The signal processing of raw local field potential data must be translated from desktop PC software algorithms to FPGA hardware, and mechanisms to resume UKF state reconstruction after seizure events will be investigated.

The UKF is a powerful tool with many possible applications in biological research and biomedical devices that are only beginning to be explored. In this paper, we have demonstrated that a model that at first appears to be too complex to be realized in a low-power system can be scaled down in a way that maintains the quality of state forecasting and reconstruction. It is hoped that the results presented here will aid others in realizing UKF systems in highly portable embedded platforms.

## Author contributions

**Conceptualization:** Bruce J. Gluckman.

**Data curation:** Philip P. Graybill.

**Formal analysis:** Philip P. Graybill.

**Funding acquisition:** Philip P. Graybill, Bruce J. Gluckman, Mehdi Kiani.

**Investigation:** Philip P. Graybill.

**Methodology:** Philip P. Graybill, Bruce J. Gluckman.

**Resources:** Bruce J. Gluckman, Mehdi Kiani.

**Software:** Philip P. Graybill.

**Supervision:** Bruce J. Gluckman, Mehdi Kiani.

**Validation:** Bruce J. Gluckman.

**Visualization:** Philip P. Graybill.

**Writing – original draft:** Philip P. Graybill, Bruce J. Gluckman.

**Writing – review & editing:** Philip P. Graybill, Bruce J. Gluckman, Mehdi Kiani.

## Footnotes

1 Before this transformation, the sigma points may be regenerated using 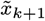 and 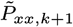. This optional step has the advantage of placing the sigma points on the isoprobability defined by 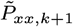 before transformation. In this paper, we have omitted this step because of the simple linear nature of our observation function.

## References

1. Bahari F, Tulyaganova C, Billard M, Alloway K, Gluckman BJ. The neural basis for sleep regulation - Data assimilation from animal to model. Conference Record - Asilomar Conference on Signals, Systems and Computers. 2017; p. 1061–1065. doi:10.1109/ACSSC.2016.7869532.

2. Voss HU, Timmer J, Kurths J. Nonlinear dynamical system identification from uncertain and indirect measurements. International Journal of Bifurcation and Chaos in Applied Sciences and Engineering. 2004;14(6):1905–1933. doi:10.1142/S0218127404010345.

3. Schiff SJ, Sauer T. Kalman filter control of a model of spatiotemporal cortical dynamics. Journal of Neural Engineering. 2008;5(1):1–8. doi:10.1088/1741-2560/5/1/001.

4. Saatci E, Akan A. Dual Unscented Kalman Filter and Its Applications to Respiratory System Modelling. In: Moreno VM, Pigazo A, editors. Kalman Filter: Recent Advances and Applications. Vienna, AUS: IntechOpen; 2009.

5. Ullah G, Schiff SJ. Assimilating Seizure Dynamics. PLoS Computational Biology. 2010;6(5). doi:10.1371/journal.pcbi.1000776.

6. Meskin N, Nounou H, Nounou M, Datta A. Parameter estimation of biological phenomena: An unscented Kalman filter approach. IEEE/ACM Transactions on Computational Biology and Bioinformatics. 2013;10(2):537–543. doi:10.1109/TCBB.2013.19.

7. Kuhlmann L, Freestone DR, Manton JH, Heyse B, Vereecke HEM, Lipping T, et al. Neural mass model-based tracking of anesthetic brain states. NeuroImage. 2016;133:438–456. doi:10.1016/j.neuroimage.2016.03.039.

8. Albers DJ, Levine M, Gluckman B, Ginsberg H, Hripcsak G, Mamykina L. Personalized glucose forecasting for type 2 diabetes using data assimilation. PLoS Computational Biology. 2017;13(4). doi:10.1371/journal.pcbi.1005232.

9. Bahari F, Kimbugwe J, Alloway KD, Gluckman BJ. Model-based analysis and forecast of sleep-wake regulatory dynamics: Tools and applications to data. Chaos. 2021;31. doi:10.1063/5.0024024.

10. Soh J, Wu X. A FPGA-based approach to attitude determination for nanosatellites. In: Proceedings of the 7th IEEE Conference on Industrial Electronics and Applications (ICIEA). IEEE; 2012. p. 1700–1704.

11. Soh J, Wu X. A scalable, portable, FPGA-based implementation of the Unscented Kalman Filter. In: NASA/ESA Conference on Adaptive Hardware and Systems (AHS); 2014. p. 127–134.

12. Soh J, Wu X. An FPGA-Based Unscented Kalman Filter for System-On-Chip Applications. IEEE Transactions on Circuits and Systems II: Express Briefs. 2017;64(4):447–451. doi:10.1109/TCSII.2016.2565730.

13. Fico VM, Arribas CP, Soaje AR, Prats MAM, Utrera SR, Vazquez ALR, et al. Implementing the unscented Kalman filter on an embedded system: A lesson learnt. In: Proceedings of the IEEE International Conference on Industrial Technology. Seville, Spain: IEEE; 2015. p. 2010–2014.

14. Valade A, Acco P, Grabolosa P, Fourniols JY. A study about Kalman filters applied to embedded sensors. Sensors (Switzerland). 2017;17(2810):1–18. doi:10.3390/s17122810.

15. Zhu X, Jiang R, Chen Y, Hu S, Wang D. FPGA implementation of Kalman filter for neural ensemble decoding of rat’s motor cortex. Neurocomputing. 2011;74(17):2906–2913. doi:10.1016/j.neucom.2011.03.044.

16. Diniz Behn CG, Booth V. Simulating Microinjection Experiments in a Novel Model of the Rat Sleep-Wake Regulatory Network. Journal of Neurophysiology. 2010;103(4):1937–1953. doi:10.1152/jn.00795.2009.

17. Julier SJ, Uhlmann JK. New extension of the Kalman filter to nonlinear systems. In: Signal Processing, Sensor Fusion, and Target Recognition VI. Orlando, FL, USA: SPIE; 1997. p. 182–193.

18. Julier S, Uhlmann J, Durrant-Whyte HF. A New Method for the Nonlinear Transformation of Means and Covariances in Filters and Estimators. IEEE Transactions on Automatic Control. 2000;45(3):477–482.

19. van der Merwe R, Wan EA. The square-root unscented Kalman filter for state and parameter-estimation. In: ICASSP, IEEE International Conference on Acoustics, Speech and Signal Processing. vol. 6. Salt Lake City, Utah, USA; 2001. p. 3461–3464.

20. Julier SJ, Uhlmann JK. Reduced sigma point filters for the propagation of means and covariances through nonlinear transformations. Proceedings of the American Control Conference. 2002;2:887–892. doi:10.1109/ACC.2002.1023128.

21. Julier SJ. The Spherical Simplex Unscented Transformation. In: Proceedings of the American Control Conference. vol. 3. Denver, Colorado, USA: IEEE; 2003. p. 2430–2434.

22. Simon D. Optimal State Estimation: Kalman, H-Infinity, and Nonlinear Approaches. Hoboken, NJ, USA: John Wiley & Sons, Inc.; 2006.

23. Sedigh-Sarvestani M, Schiff SJ, Gluckman BJ. Reconstructing Mammalian Sleep Dynamics with Data Assimilation. PLoS Computational Biology. 2012;8(11). doi:10.1371/journal.pcbi.1002788.

24. Graybill P, Gluckman BJ, Kiani M. Toward a Wearable Data Assimilation Platform. In: IEEE Biomedical Circuits and Systems Conference. Nara, Japan; 2019.

25. Billard MW, Bahari F, Kimbugwe J, Alloway KD, Bruce J. The systemDrive : a Multisite, Multiregion Microdrive with Independent Drive Axis Angling for Chronic Multimodal Systems Neuroscience Recordings in Freely Behaving Animals. eNeuro. 2018;5(6):1–19.

26. Rhudy M, Gu Y, Gross J, Napolitano MR. Evaluation of matrix square root operations for UKF within a UAV GPS/INS sensor fusion application. International Journal of Navigation and Observation. 2011;2011. doi:10.1155/2011/416828.

27. Straka O, Duník J, Šimandl M, Havlík J. Aspects and Comparison of Matrix Decompositions in Unscented Kalman Filter. In: American Control Conference. Washington, DC, USA: IEEE; 2013. p. 3075–3080.

28. Sakai A, Kuroda Y. Discriminatively Trained Unscented Kalman Filter for Mobile Robot Localization. Journal of Advanced Research in Mechanical Engineering. 2010;1(3):153–161.

29. Duník J, Šimandl M, Straka O. Unscented Kalman Filter: Aspects and Adaptive Setting of Scaling Parameter. IEEE Transactions on Automatic Control. 2012;57(9):2411–2416.

30. Straka O, Duník J, Šimandl M. Unscented Kalman filter with advanced adaptation of scaling parameter. Automatica. 2014;50(10):2657–2664. doi:10.1016/j.automatica.2014.08.030.

31. Scardua LA, da Cruz JJ. Complete offline tuning of the unscented Kalman filter. Automatica. 2017;80:54–61. doi:10.1016/j.automatica.2017.01.008.

32. Turner R, Rasmussen CE. Model based learning of sigma points in unscented Kalman filtering. Neurocomputing. 2012;80:47–53. doi:10.1016/j.neucom.2011.07.029.

33. Duník J, Straka O, Kost O, Havlík J. Noise covariance matrices in state-space models: A survey and comparison of estimation methods—Part I. International Journal of Adaptive Control and Signal Processing. 2017;31:1505–1543. doi:10.1002/acs.2783.

34. Vachhani P, Narasimhan S, Rengaswamy R. Robust and reliable estimation via Unscented Recursive Nonlinear Dynamic Data Reconciliation. Journal of Process Control. 2006;16(10):1075–1086. doi:10.1016/j.jprocont.2006.07.002.

35. Narasimhan S, Rengaswamy R. Reply to Comments on “Robust and reliable estimation via unscented recursive nonlinear dynamic data reconciliation” (URNDDR). Journal of Process Control. 2009;19(4):719–721. doi:10.1016/j.jprocont.2008.08.002.

36. Kolås S, Foss BA, Schei TS. Constrained nonlinear state estimation based on the UKF approach. Computers and Chemical Engineering. 2009;33(8):1386–1401. doi:10.1016/j.compchemeng.2009.01.012.

37. Simon DJ. Kalman Filtering With State Constraints: A Survey of Linear and Nonlinear Algorithms. IET Control Theory & Applications. 2010;4(8):1303–1318.

38. Teixeira BOS, Tôrres LAB, Aguirre LA, Bernstein DS. On unscented Kalman filtering with state interval constraints. Journal of Process Control. 2010;20(1):45–57. doi:10.1016/j.jprocont.2009.10.007.

